# Assessing the fractional contributions of static, slow and fast dynamic scatterer components to the flow index derived by continuous wave diffuse correlation spectroscopy

**DOI:** 10.64898/2026.08.14.744820

**Authors:** Neda Mogharari, Michał Kacprzak, Dawid Borycki

## Abstract

Continuous wave diffuse correlation spectroscopy (cw-DCS) is a noninvasive optical technique to monitor the tissues’ blood flow changes. This technique measures the tissue blood flow index (BFI) by evaluating the decay rate of the autocorrelation function. The derived BFI is proportional to mean squared displacements of the red blood cells considered as the fast-dynamic scatterer component of tissue in time. However, biological tissue contains static scatterer component and slow-dynamic scatterer component which affect the decay rate of autocorrelation function and as a result the derived BFI. In this study, we assessed the fractional contribution of static, slow-dynamic and fast-dynamic scatterer components of a medium in the flow index derived by cw-DCS. The measurements performed on Agar-based phantom with tube showed that presence of static scatterer component and slow-dynamic scatterer component led to substantial underestimation (≈ 123%) of the flow index derived by Siegert relation, compared to effective diffusion coefficient of fast-dynamic scatterers components derived by modified Siegert relation and bi-exponential model. The less underestimation was observed for the corresponding parameters obtained from the liquid phantom measurements (≈ 25%) as well as during the forearm occlusion test and respiratory challenges (≈ 16% − 26%).

## 1. Introduction

Adequate blood flow is essential for maintaining tissue viability by continuously delivering oxygen and nutrients while removing the metabolic waste products. Therefore, accurate monitoring of tissue and cerebral blood flow (CBF) is of considerable importance for the early detection and assessment of disorders associated with impaired perfusion and ischemia [1]. Numerous noninvasive techniques have been developed to monitor blood flow; however, each possesses limitations that restrict its widespread clinical application. Transcranial Doppler ultrasound provides measurements of blood flow velocity in major arteries but does not directly assess microvascular perfusion [2]. Laser Doppler flowmetry primarily interrogates superficial tissues and generally requires direct optical access to the tissue of interest [3]. Positron emission tomography (PET) is considered one of the reference methods for quantitative CBF assessment, yet its dependence on radioactive tracers limits repeated examinations and continuous bedside monitoring [4]. Magnetic resonance imaging (MRI)-based perfusion methods offer high spatial resolution and detailed anatomical information but suffer from limited temporal resolution and are unsuitable for continuous monitoring [5]. Furthermore, both PET and MRI require costly, non-portable instrumentation and specialized imaging facilities, making their routine use for bedside or long-term blood flow monitoring challenging.

Diffuse correlation spectroscopy (DCS) is an optical technique to monitor tissue and CBF changes. Due to noninvasiveness, portability, and relatively low cost, this technique has been widely used in neuroimaging and tissue blood flow studies [6]. Blood flow monitoring using DCS has been validated against a variety of established imaging modalities. These include color-weighted power Doppler ultrasound in murine tumor models [7]; laser Doppler flowmetry in rats [8]; xenon-enhanced computed tomography in humans [9]; phase-encoded velocity-mapping MRI during hypercapnia in children [10]; arterial spin labeling MRI during muscle occlusion in humans [11], and during hypercapnic challenges on rodents [12] and healthy adults [13]; as well as blood oxygenation level–dependent MRI in the rat brain during electrical forepaw stimulation [14] and human brain during respiratory challenges [15].

In continuous wave DCS (cw-DCS), a continuous wave long coherent near infrared light source is used to illuminate the tissue. The temporal light intensity fluctuation of the near-infrared light diffused through the tissue is used to calculate the autocorrelation function. The temporal decay rate of the autocorrelation function is subsequently quantified to estimate the tissue blood flow changes [16]. The photon scattering from both static and dynamic scatterer components of tissue contribute to the decorrelation of the laser light and decay of autocorrelation function. The blood flow index (BFI) measured by DCS is proportional to the mean squared displacement of the red blood cells (RBCs) [17] as fast-dynamic scatterer component of tissue. However, static scatterer component such as extravascular structure of tissue (e.g., nuclei, mitochondria) [18], and slow-dynamic scatterer component such as intracellular structure of tissue with slower motility [19] can also affect the decay of the autocorrelation function. This conceptual overview has been presented in figure 1.

**Figure 1.**
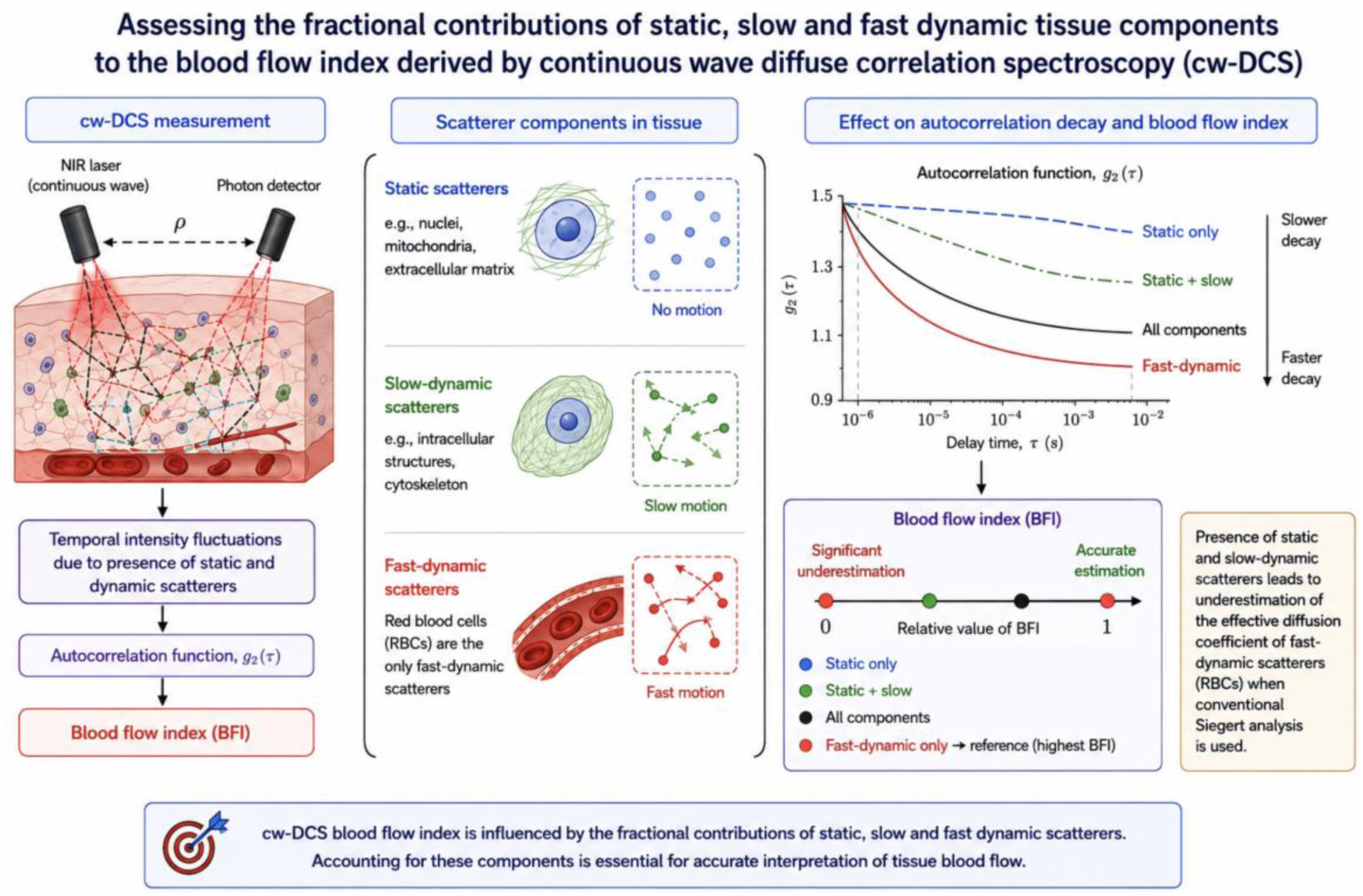
Conceptual overview of the contributions of static, slow-dynamic, and fast-dynamic scatterer components to the BFI (blood flow index) measured by cw-DCS (continuous-wave diffuse correlation spectroscopy). Tissue contains static scatterers (e.g., extracellular structures), slow-dynamic scatterers (e.g., motility of intracellular components), and fast-dynamic scatterers (RBCs). In cw-DCS, temporal intensity fluctuations of diffused scattered coherent light are analyzed through the *g*_2_(*τ*) (intensity autocorrelation function), to estimate the BFI. Because *g*_2_(*τ*) reflects the combined contributions of all scatterer components, analysis based on the Siegert relation attributes part of the decorrelation arising from static and slow-dynamic tissue components to the blood flow, resulting in an underestimation of the absolute blood flow index measured by cw-DCS. The modified Siegert relation and bi-exponential models presented in this work can distinguish these contributions, enabling estimation of the effective diffusion coefficient associated with fast-dynamic scatterers while quantifying the relative contributions of static and slow-dynamic scatterer tissue components to the estimated cw-DCS derived blood flow index.

In this study, we conducted series of measurements on Agar-based phantom with tube and liquid phantoms as well as muscle occlusion test and respiratory challenges. Then we performed three data analyses approaches including Siegert relation, modified Siegert relation and bi-exponential models and compared their derived flow related parameters. These analyses help to better understand the fractional contribution of static, slow-dynamic and fast-dynamic scatterer components of medium to the flow index measured by cw-DCS, especially when the absolute blood flow index is required.

## 2. Theory

The flow index measured by cw-DCS is based on the evaluation of the temporal autocorrelation function of the light intensity fluctuations caused by scatterers and displacements of RBCs. To assess the dynamic of scatterers and RBCs, the normalized intensity autocorrelation function *g*_2_(*τ*) is calculated by [20]:

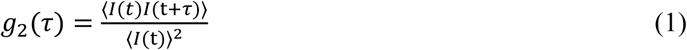

where t is the absolute arrival time of photons, *I* is the light intensity fluctuations of detected photons, and *τ* is the time of scatterers’ displacement (the lag time).

In ergodic samples, in which all scatterers are moving, and the medium does not contain any static scatterer, the intensity autocorrelation function *g*_2_(*τ*) is related to the normalized electric field autocorrelation function *g*_1_(*τ*) by the Siegert relation [20]:

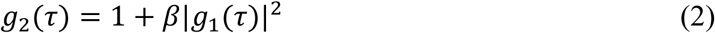

where *β* is the coherence parameter. In the case of using a long coherence length laser, photon detection with a single mode fiber gives *β* = 0.5 for the unpolarized light [21]. The normalized electric field autocorrelation function *g*_1_(*τ*) can be expressed by [22]:

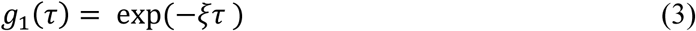

The electric field autocorrelation function decay rate *ξ* can be calculated as follows [23]:

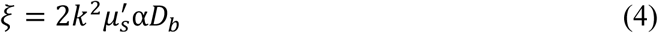

where 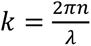 is the wavenumber in the medium with refractive index of *n* and light wavelength of *λ*, *μ*′s is the reduced scattering coefficient, and *D_b_* is the effective diffusion coefficient. The product of α*D_b_* which is proportional to mean squared displacement of the scatterers in time, reflects the BFI in the tissue. In *αD*_b_ term, the *α* refers to the fraction of dynamic scatterers to total scatterers (ranging from 0 to 1) [24].

As, biological tissues include both static scatterer (e.g., extravascular structure of tissue) [18] and dynamic scatterer (e.g., intracellular structure of tissue with slower motility, moving RBCs) [19] components, they are not ergodic samples. Therefore, the contribution of static scatterer components should be excluded from the autocorrelation decay rate. For a not ergodic medium, the modified Siegert relationship is as follows [25]:

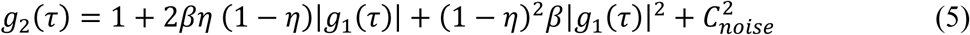

where *η* is the ratio of the averaged light intensity fluctuation of the static scatterers to total scatterers (static and dynamic) which varies from 0 to 1. The last term 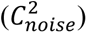 represents the experimental noise. To estimate the *αD*_b_ of dynamic scatterer components (*αD*_b (Dynamic)_) while excluding the contributions of static scatterer components, equation (5) was incorporated into equation (3) and (4).

In non-ergodic mediums, to evaluate the contribution of slow-dynamic (e.g., intracellular structure of tissue with slower motility) and fast-dynamic (moving RBCs) [19] scatterer components on the cw-DCS derived flow index, the *g*_1_(*τ*) can be calculated using a bi-exponential model [26]:

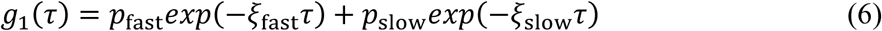

where *ξ*_fast_ refers to decay rate caused by fast-dynamic scatterer component. *p*_fast_ represents the fractional contribution of the fast-dynamic scatterer component with decay rate of *ξ*_fast_.

*ξ*_slow_ corresponds to decay rate of slow-dynamic scatterer components. The *p*_slow_ which is the fractional contribution of the slow-dynamic scatterer component with decay rate of *ξ*_slow_, can be derived as 1 − *p*_fast_. The *p*_fast_ and *p*_slow_ varies from 0 to 1. To distinguish the effective diffusion coefficient of slow-dynamic scatterers components (*D*_slow_) and effective diffusion coefficient of fast-dynamic scatterers components (*D*_fast_), equation (6) can be incorporated into equations (4) and (5).

The bi-exponential model to estimate the fractional contribution of the tissues’ slow and fast components has been also introduced in intravoxel incoherent motion magnetic resonance imaging (IVIM-MRI) technique. In this technique the resembled of *D*_slow_ and *D*_fast_ parameters originate from water molecules diffusion within the extravascular structure of tissue and blood perfusion within the microvascular structure of tissue. Moreover, the parameter similar as *p*_fast_, represent the volume fraction of the blood flow within the tissues’ microvessels [27, 28].

## 3. Methods

### 3.1 Instrumentation

The detailed description of cw-DCS system used to acquire the experimental data of this study was previously reported in our works [Submitted]. The optical setup is equipped with a picosecond pulsed laser (VisIR-765-HP "STED", PicoQuant, Germany) at wavelength of 765 nm using 80 MHz repetition rate and a continuous-wave laser (DL785-100-SO, Crystalaser, USA) at wavelength of 785 nm with coherence length > 10 m as sources to measure the optical properties and dynamic properties, respectively. Two single-photon avalanche diodes (PDM, Micro Photon Devices), and two time-correlated single photon counting boards (SPC-150, Becker&Hickl, Germany) were synchronized with the pulsed laser. The SPCM data acquisition software (Multifunctional 64-bit Data Acquisition Software, Becker & Hickl) [29] was used to record the signals. The distance between the source and detection fiber tips was 2 cm.

### 3.2 Physical phantoms

To evaluate the contribution of static, slow-dynamic, and fast-dynamic scatterer components of tissue, two phantoms were designed.

#### Agar-based phantom with tube

The detailed description of the preparation of Agar-based phantom with tube has been explained in our previous work [Submitted]. We utilized a container, 3D printed from black ABS material, with parallel holes to pass transparent silicone tubes. The container was filled with an Agar mixture as static and slow-dynamic scatterers medium. The SMOFlipid 20 (Fresenius Kabi) was added to the mixture to obtain a turbid medium with reduced scattering coefficient of *μ*′s = 10 cm^-1^ and absorption coefficient of *μ*a = 0.03 cm^-1^. The silicone tube with the inner diameter of 0.1 cm, was connected to a programmable syringe pump (NE-1000 Programmable Single Syringe Pump), which enables generating the desired pump flow rates (*F*_pump_) of fast-dynamic scatterers. The liquid mixture to pump through the tubes was also prepared using SMOFlipid as the scatterers (i.e., mimicking the scattering properties of RBCs) and distilled water. The measured optical properties of this mixture were *μ*′s = 10 cm^-1^ and *μ*a = 0.03 cm^-1^ and were prepared according to the recipe described in [30, 31]. The source and detector fibers were confined in a 3D printed holder and placed at the surface of the container. The distance between the rows of the tubes and the optical fibers tip was 0.5 cm.

### Liquid phantom

A one-layer 3D container was designed from the black ABS material. The mixtures to fill the container included SMOFlipid as scatterer, distilled water and Glycerol (Vegetable Glycerin 99.5% pharmaceutical grade, Bioswena, Poland) to slow down the dynamic of scatterers. Three mixtures with 40% glycerol concentration, 20% glycerol concentration and without glycerol (0% glycerol concentration) were prepared. To adjust the desired reduced scattering coefficient, different concentrations of SMOFlipid were added to each mixture according to the recipe described in [30, 31]. The optical properties of all mixtures, were *μ*a = 0.03 cm^-1^ and *μ*′s = 10 cm^-1^. Same as previous phantom, the source and detector fibers were confined in a 3D printed holder and placed at the surface of the container.

The absorption property of the sample doesn’t substantially affect the scatterers displacements [32]. Therefore, no absorber was added to the mixtures of each Agar-based phantom with tube and liquid phantoms.

### 3.3 In-vivo measurements protocol

The study was approved by the Ethics Committee of the Military Medical Chamber in Warsaw, Poland (Approval No. 17/23). All participants provided written informed consent before participation. The study was conducted in accordance with the principles of the Declaration of Helsinki.

#### Forearm occlusion

The forearm occlusion trial involved placing a cuff pressure (tourniquet) around the upper arm, inflating it to occlude the blood flow for a certain duration, then releasing it. The occlusion trial started with 40 s of baseline, followed by occlusion pressures of 60, 100, 140 and 180 mmHg which each lasted 40s and ended with 40 s of release. The occlusion trial was repeated twice.

#### Respiratory challenges

The detailed description of the respiratory challenges was reported in our previous work [15]. To change the CBF, two respiratory challenges of breath hold (BH) and Hyperventilation (HV) were performed which caused an increase and reduction of the CBF, respectively. The BH trial consisted of 90 s long period of self-paced breathing, followed by 30 s of breath-holding and ended with 90 s of self-paced breathing. The HV trial consisted of an 80 s long period of self-paced breathing, continued with 40 s of fast-breathing and ended by 80 s of self-paced breathing. Each BH and HV trial was repeated two times.

#### Volunteers

One (female) and eight (four females and four males) healthy adult volunteers participated in forearm occlusion and respiratory challenges, respectively. As the forearm occlusion was performed by a single subject, the related observations should be interpreted as a proof-of-concept measurement.

### 3.4 Data analyses

#### Optical properties

The measured distribution of times of flight (DTOF) of each acquired data was used to calculate the optical properties of *μ_a_* and 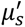, based on statistical moment analysis of the DTOF curve [33]. In case of in-vivo tests, although the tissue’s optical properties vary between baseline and muscle occlusion states as well as baseline and respiratory challenges, previous studies have shown that minor variations in *μ*ₛ′ have a negligible effect on the derived *αD_b_* and BFI [32]. Therefore, we used the same *μ*ₐ and *μ*ₛ′ measured during the baseline for all states.

#### Fitting ranges to calculate the autocorrelation function

For both phantom and in-vivo measurements, the autocorrelation functions *g*_2_(*τ*) were obtained for time range of 10^−6^to 10^−2^ s. This lag time range was chosen to capture the full decorrelation caused by static, slow-dynamic and fast-dynamic scatterer components, while maintaining sufficient signal-to-noise ratio. Then to calculate the *g*_1_(*τ*), the lag time ranges were chosen, where the slowest decay curve *g*_2_(*τ*) measured on phantoms and during in-vivo tests is *g*_2_(*τ*) = 1 (figure 2, red curves). These lag ranges were then used for fitting other *g*_1_(*τ*) curves.

**Figure 2.**
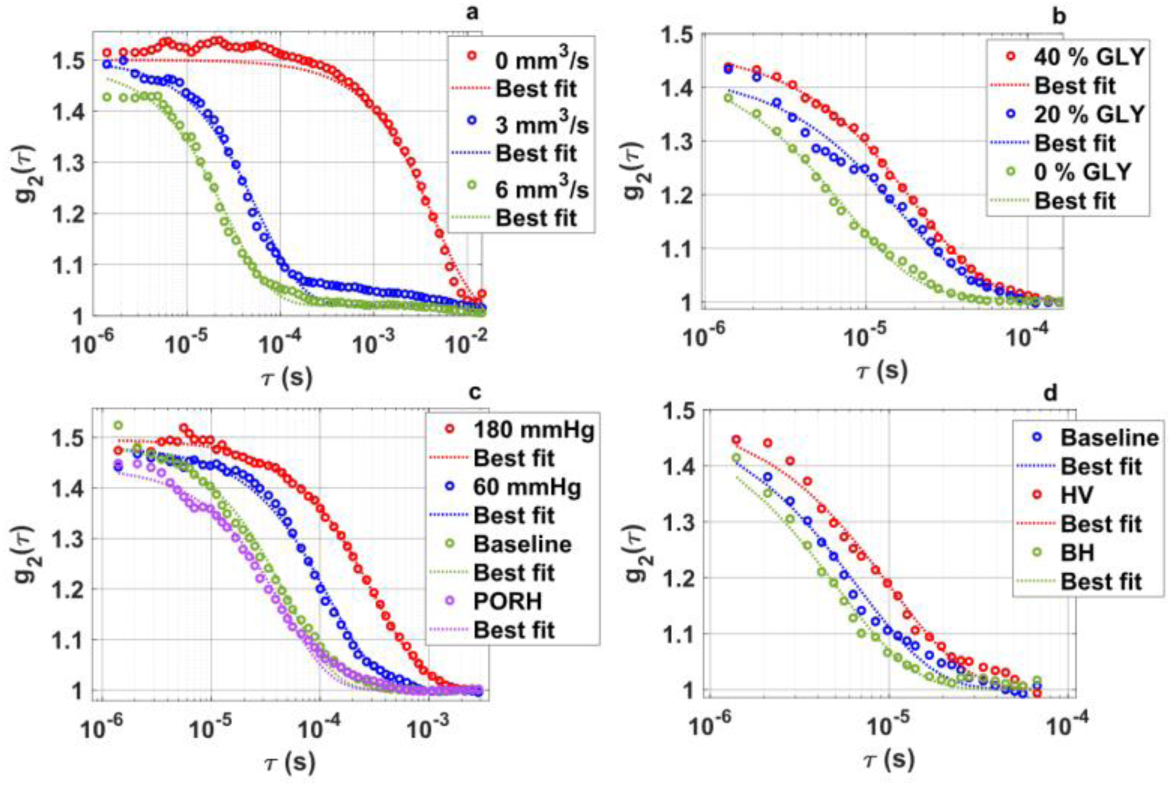
The fitting range of autocorrelation function *g*_2_(τ) to calculate the *αD_b_* of a) Agar-based phantom with tube, and b) Liquid phantoms, as well as BFI during c) forearm occlusion, and d) respiratory challenges.

#### Brownian model

At low flow rates within the tube of the tissue-mimicking phantom connected to the syringe pump, the Brownian like motion of scatterers and RBCs is dominant in the decorrelation of the signal detected by DCS [34, 35]. Therefore, for both tissue mimicking phantom and muscle occlusion data analysis, the *αD_b_* was calculated using Brownian model. The lsqcurvefit function in MATLAB was used to fit the obtained *g*_1_(*τ*) from theory to the measured *g*_1_(*τ*).

#### Integration time

In both phantom and in-vivo data analyses, the *g*_2_(*τ*) was obtained with 3s integration time.

## 4. Results

*Agar-based phantom with tube*: Figure 3 shows the *αD_b_* derived using Siegert relation and *αD_b_*_(Dynamic)_ and *η* derived using modified Siegert relation of the data measured on Agar-based phantom with tube. The mean *αD_b_*_(Dynamic)_ calculated at *F*_pump_ from 1 to 6 mm^3^/s on average is 49% higher than *αD_b_*. Due to solid-like structure of the Agar, the *η* is high (close to 1), especially at baseline (*F*_pump_ = 0). While, increasing the *F*_pump_ led to relatively decrease the *η* in comparison to baseline.

**Figure 3.**
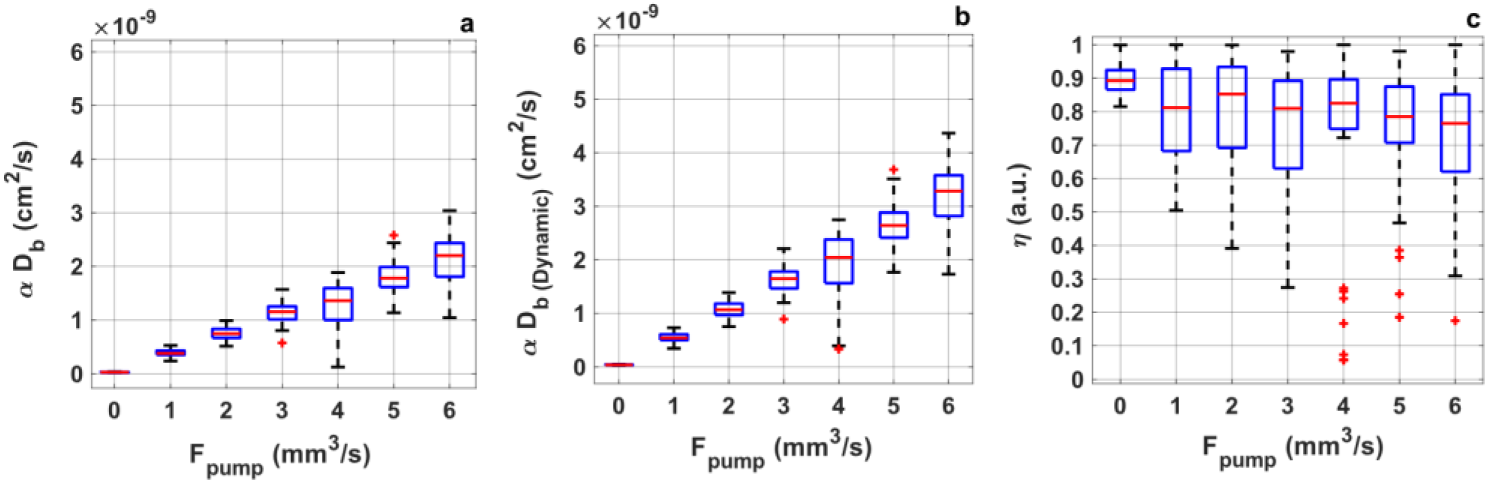
The flow parameters obtained from agar-based phantom with tube. a) αD_b_ derived by Siegert relation, as well as b) αD_b(Dynamic)_, and c) η derived by modified Siegert relation of the data measured on Agar-based phantom with tube at baseline and six *F*_pump_ (pump flow rate). Each boxplot was generated from 100 data points. In each boxplot, the central line represents the median, while the bottom and top edges are the 25-th and 75-th percentiles respectively. The whiskers extend to the most extreme data points and the plus markers are considered outliers.

Figure 4 shows the *p*_fast_, *p*_slow_, *D*_fast_ and *D*_slow_ derived by bi-exponential models measured on Agar-based phantom with tube. At baseline (*F*_pump_ = 0), the *p_fast_* is around 0.5. Increasing the *F*_pump_ led to increase the mean value of this parameter to around 0.8 and remained relatively constant at all *F*_pump_ with some minor changes. The *p*_slow_, at baseline is substantially higher than this value at all *F*_pump_. As expected, increasing the pump flow, increased the *D*_fast_; while, *D*_slow_ remained relatively constant with minor changes. The mean *D*_fast_ calculated at *F*_pump_ from 1 to 6 mm^3^/s on average is 52% and 123% higher than *αD_b_*_(Dynamic)_ and *αD_b_*, respectively. The estimated *D*_fast_ is relatively two orders of magnitude higher than *D*_slow_.

**Figure 4.**
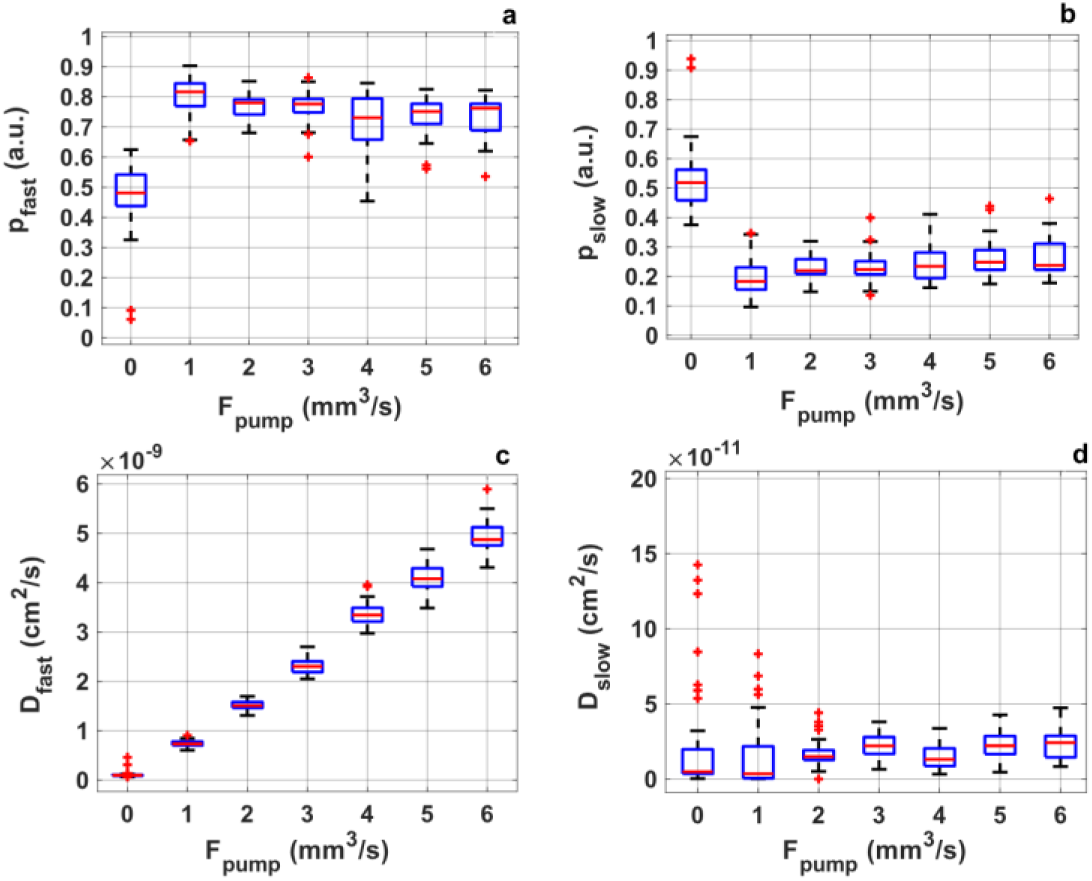
The flow parameters obtained from agar-based phantom with tube. a) *p*_fast_ (fractional contribution of the fast-dynamic scatterer component), b) *p*_slow_ (fractional contribution of the slow-dynamic scatterer component), c) *D*_fast_ (effective diffusion coefficient of fast-dynamic scatterers components), and d) *D*_slow_ (effective diffusion coefficient of slow-dynamic scatterers components) derived by bi-exponential model measured on Agar-based phantom with tube at baseline and six *F*_pump_ (pump flow rate). Each boxplot was generated from 100 data points. In each boxplot, the central line represents the median, while the bottom and top edges are the 25-th and 75-th percentiles respectively. The whiskers extend to the most extreme data points and the plus markers are considered outliers.

### Liquid phantoms

Figure 5 shows the *αD_b_* derived by Siegert relation as well as the *αD_b_*_(*Dynamic*)_ and *η* derived by modified Siegert relation of the data measured on three liquid phantoms with 40%, 20% and 0% Glycerol concentration. The *αD_b_*_(Dynamic)_ averaged over all phantoms is 17% higher than the corresponding calculated *αD_b_*. The calculated *η* are close to zero, especially in phantom with 0% Glycerol concentration.

**Figure 5.**
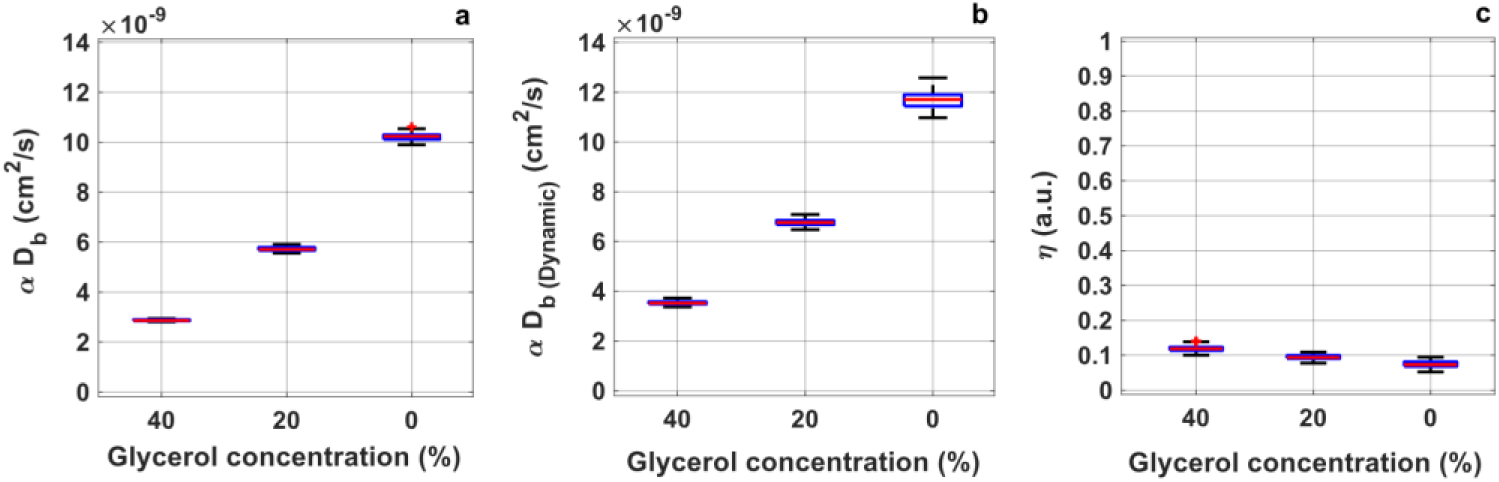
The flow parameters obtained from liquid phantoms. a) αD_b_ derived by Siegert relation as well as the b) αD_b(Dynamic)_ and c) η derived by modified Siegert relation measured on liquid phantoms. The glycerol concentrations of the phantoms are 40, 20 and 0%. Each boxplot was generated from 100 data points. In each boxplot, the central line represents the median, while the bottom and top edges are the 25-th and 75-th percentiles respectively. The whiskers extend to the most extreme data points and the plus markers are considered outliers.

As shown in figure 6, the phantoms with Glycerol (40% and 20%), have higher *p_slow_* and *D*_slow_ in comparison to phantom without Glycerol (0%). In contrast, the phantom without Glycerol (0%) show higher *p*_fast_ and *D*_fast_ in comparison to phantom with Glycerol (40% and 20%). The mean *D*_fast_ calculated from the data acquired on all three phantoms on average is 6% and 25% higher than *αD_b_*_(Dynamic)_ and *αD_b_*, respectively. The estimated *D*_fast_ of phantom with Glycerol (40% and 20%) and without Glycerol is relatively an order of magnitude and two orders of magnitude higher than *D*_slow_, respectively.

**Figure 6.**
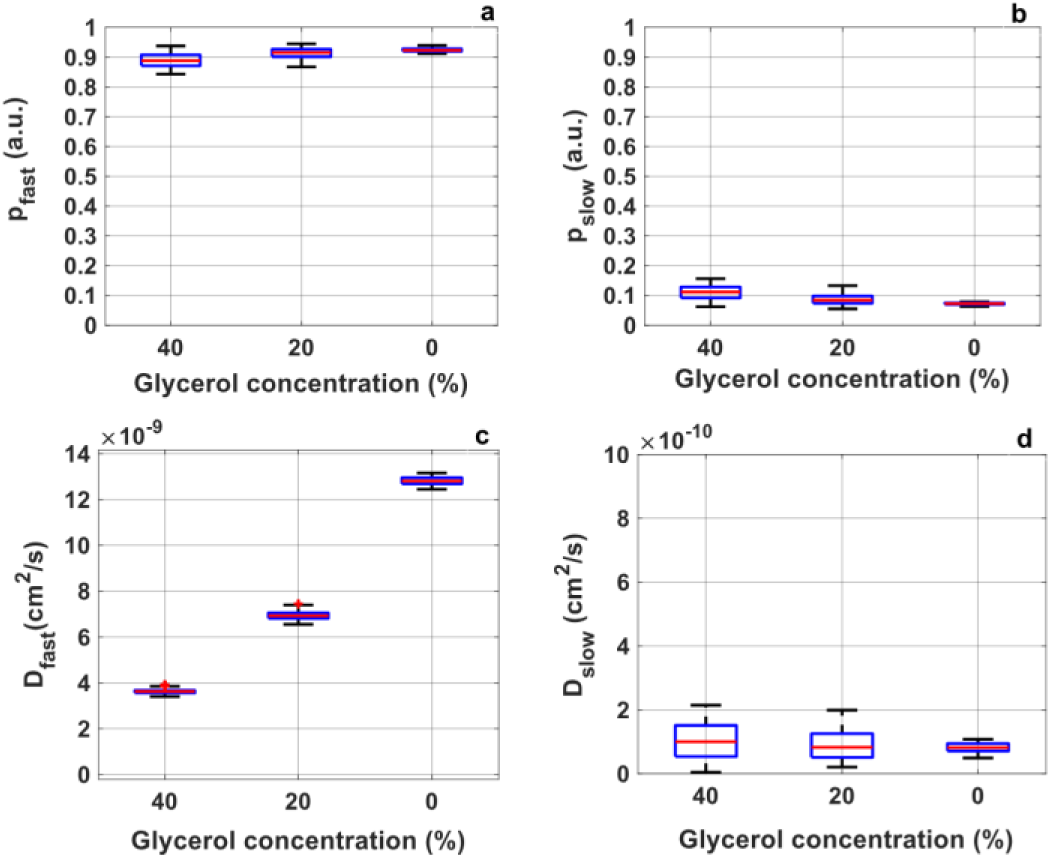
The flow parameters obtained from liquid phantoms. a) *p*_fast_ (fractional contribution of the fast-dynamic scatterer component), b) *p*_slow_ (fractional contribution of the slow-dynamic scatterer component), c) *D*_fast_ (effective diffusion coefficient of fast-dynamic scatterers components), and d) *D*_slow_ (effective diffusion coefficient of slow-dynamic scatterers components) derived by bi-exponential model measured on liquid phantoms. The glycerol concentrations of phantoms are 40, 20 and 0%. Each boxplot was generated from 100 data points. In each boxplot, the central line represents the median, while the bottom and top edges are the 25-th and 75-th percentiles respectively. The whiskers extend to the most extreme data points and the plus markers are considered outliers.

### Forearm occlusion

Figure 7 shows the BFI derived by Siegert relation as well as the *BFI*_Dynamic_ and *η* derived by modified Siegert relation from the data of forearm occlusion test. In response to occlusions, the decrease in blood flow is expected which can be seen in both BFI and *BFI*_Dynamic_. This decrease is more substantial at higher occlusion pressures (140 and 180 mmHg). The calculated *BFI*_Dynamic_ averaged over whole trial’s period including baseline, occlusion and release is 9% higher than BFI. In response to occlusion, η increased at all pressures, with higher magnitude at 140 and 180 mmHg.

**Figure 7.**
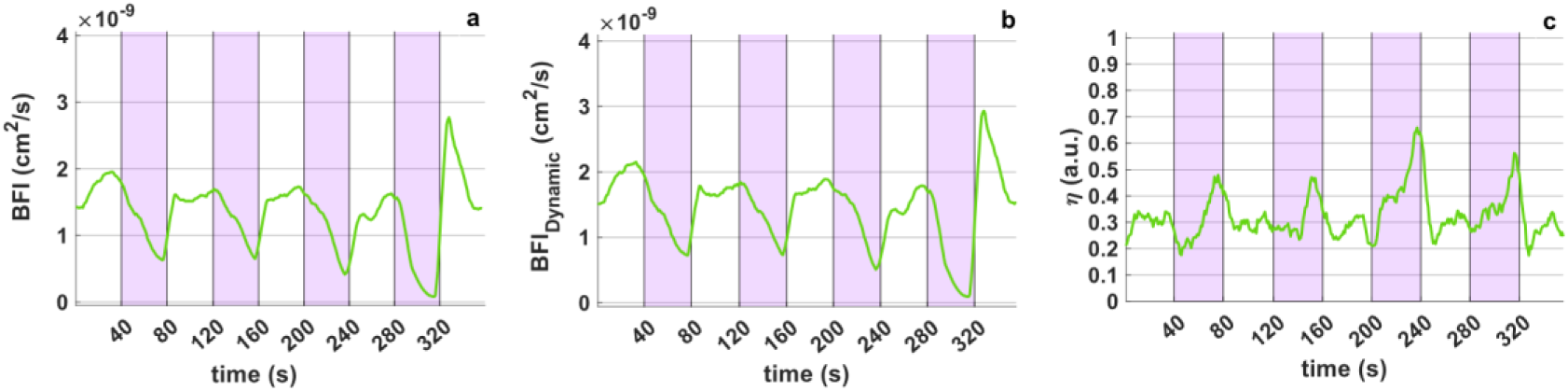
The flow parameters obtained from the performed forearm occlusion. a) BFI derived by Siegert relation and b) BFI_(Dynamic)_ and c) η derived by modified Siegert relation measured during forearm occlusion test. The green plot has been averaged from two data sets measured on one subject. The vertical purple regions indicate the occlusion periods corresponding to cuff pressures of 60, 100, 140, and 180 mmHg, from left to right.

Figure 8 shows the *p*_fast_, *D*_fast_, *p*_slow_, and *D*_slow_ calculated using bi-exponential model from the data measured during forearm occlusion trial. The *p*_fast_ shows substantial decrease in response to occlusions. In contrast, the *p*_slow_ increased in response to occlusion. The additional spikes observed in *D*_fast_ especially at 60 and 100 mmHg occlusion pressures are attributed to the reduced stability of the bi-exponential fitting procedure in estimating the parameters. The *D*_fast_ averaged over whole trial is 15% and 26% higher than *BFI*_Dynamic_ and BFI, respectively. *D*_slow_ changes in response to 60 and 100 mmHg are not substantial. However, the changes are more prominent in response to 140 and 180 mmHg pressures. The estimated *D*_fast_ is relatively an order of magnitude higher than *D*_slow_.

**Figure 8.**
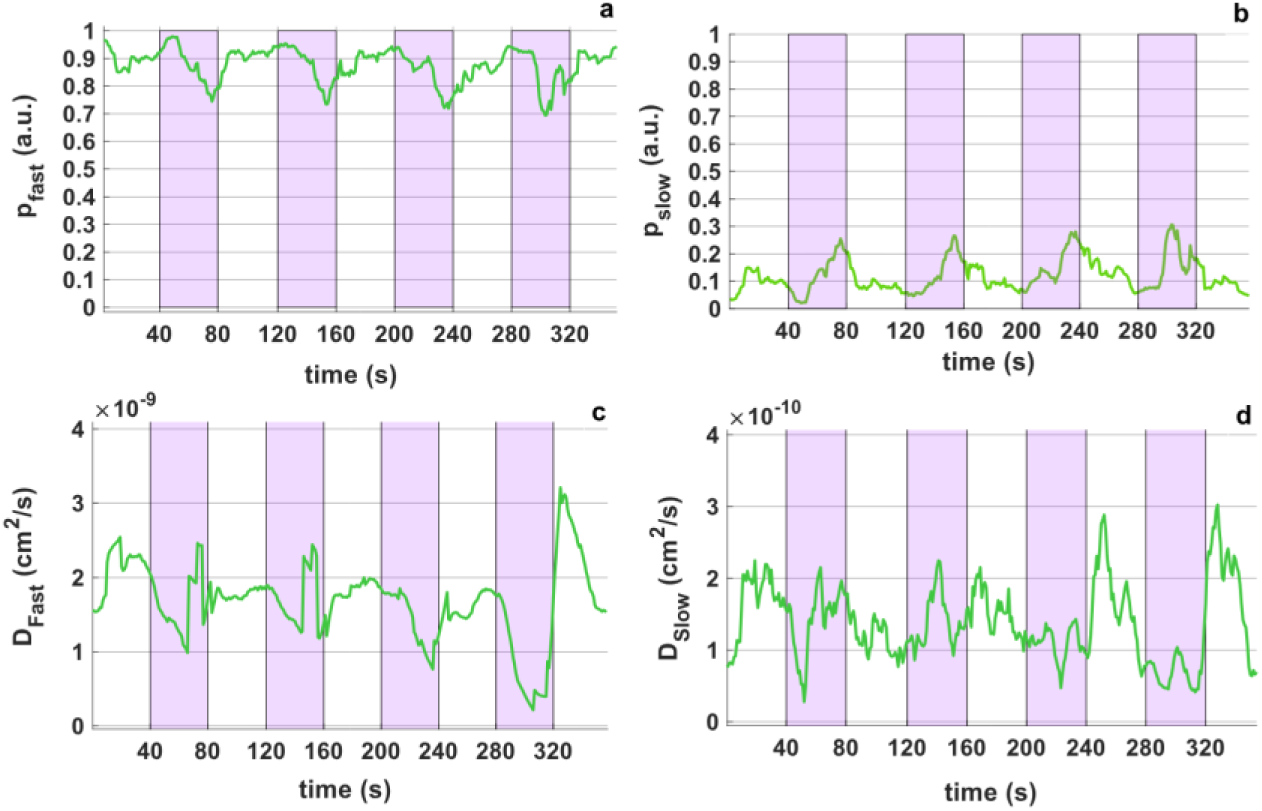
The flow parameters obtained from the performed forearm occlusion. a) *p*_fast_ (fractional contribution of the fast-dynamic scatterer component), b) *p*_slow_ (fractional contribution of the slow-dynamic scatterer component), c) *D*_fast_ (effective diffusion coefficient of fast-dynamic scatterers components), and d) *D*_slow_ (effective diffusion coefficient of slow-dynamic scatterers components) derived by bi-exponential model measured during forearm occlusion test. The green plot has been averaged from two data sets measured on one subject. The vertical purple regions indicate the occlusion periods corresponding to cuff pressures of 60, 100, 140, and 180 mmHg, from left to right.

### Respiratory challenges

To evaluate the fractional contribution of static, slow-dynamic, and fast-dynamic scatterer tissue components on the measured CBF changes by cw-DCS, BH and HV respiratory challenges were performed. Figure 9 shows the BFI calculated using Siegert relation and *BFI*_Dynamic_ and *η* calculated using modified Siegert relation. During BH, depletion of oxygen in the blood causes an increase in CBF and DCS-derived blood flow index. In contrast, in response to HV, which increases the blood oxygen level, CBF and DCS-derived blood flow index is expected to decrease [36]. The mean of *BFI*_Dynamic_ measured during whole trials’ period including baseline, BH and HV, is 3% and 7% higher than BFI, respectively. The estimated *η* didn’t change substantially in response to respiratory challenges.

**Figure 9.**
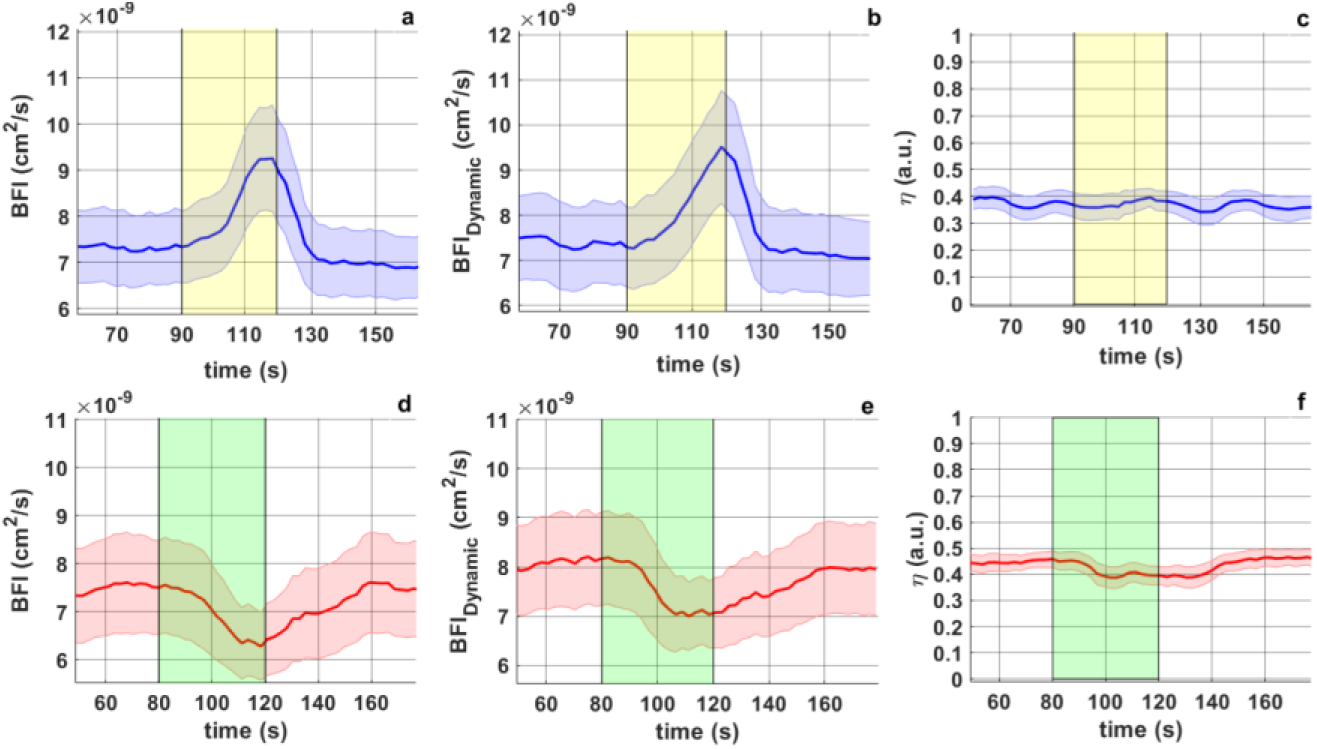
The flow parameters obtained from the performed respiratory challenges. a, d) BFI derived by Siegert relation, as well as b, e) BFI_(Dynamic)_ and c, f) η derived by modified Siegert relation measured during a, b, c) BH and d, e, f) HV respiratory challenges. The mean values (solid lines) and corresponding standard deviations (shaded areas) were obtained over 16 datasets measured on eight individuals. The vertical yellow and green regions show the BH and HV periods, respectively.

Figure 10 shows the *p*_fast_, *D*_fast_, *p*_slow_, and *D*_slow_ calculated using bi-exponential model from the data measured during respiratory challenges. The *p*_fast_ is substantially higher than *p*_slow_. Moreover, no substantial differences were observed in the both *p*_fast_ and *p*_slow_ in response to BH and HV comparing to baseline. The mean *D*_fast_ in response to BH, is 13% and 16% higher than *BFI*_Dynamic_ and BFI, respectively. The mean *D*_fast_ in response to HV, is 15% and 24% higher than *BFI*_Dynamic_ and BFI, respectively. No substantial changes in *D*_slow_, despite fluctuations were observed during the respiratory challenges. The estimated *D*_fast_ is relatively an order of magnitude higher than *D*_slow_.

**Figure 10.**
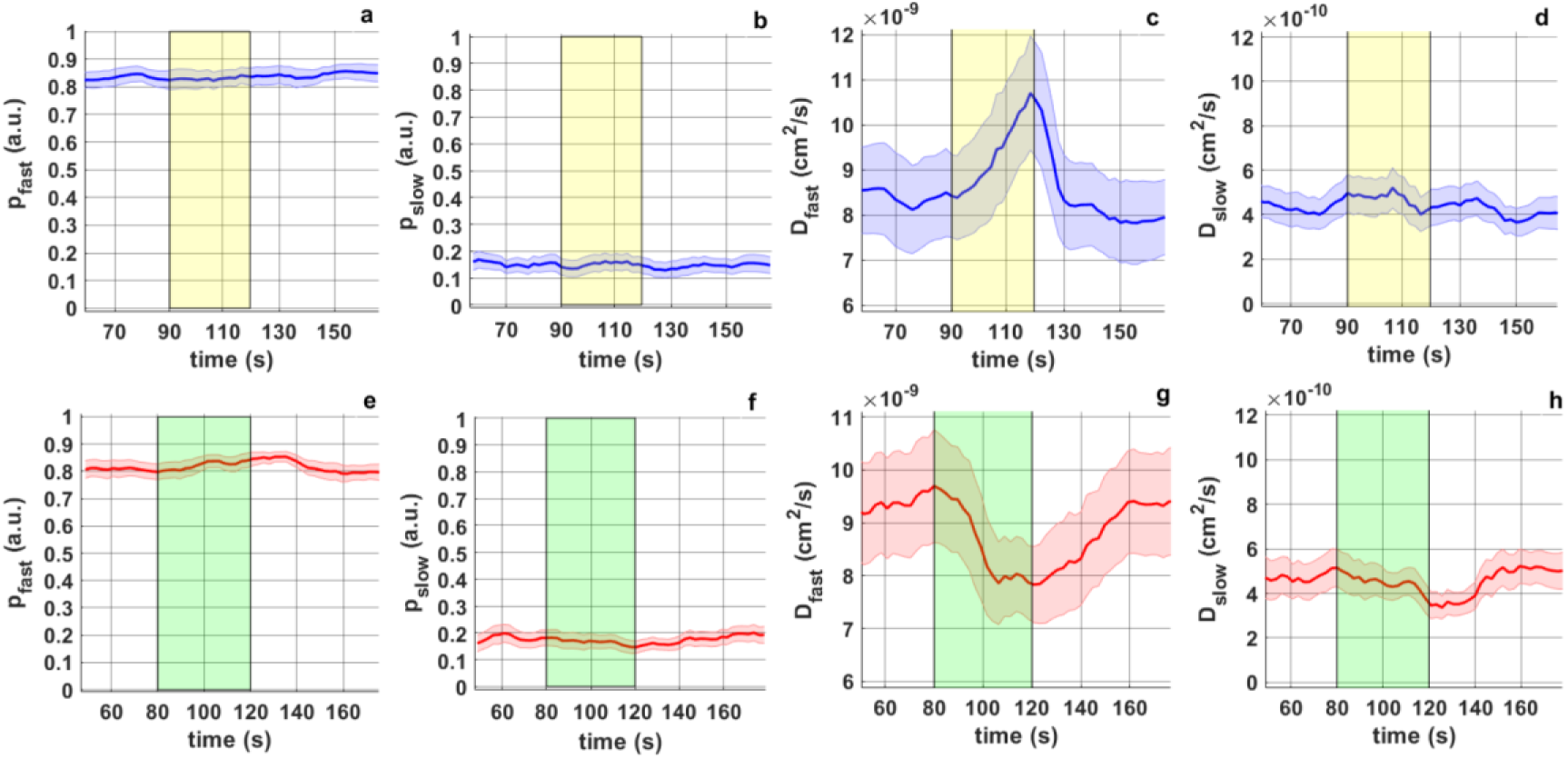
The flow parameters obtained from the performed respiratory challenges. a, e) *p*_fast_ (fractional contribution of the fast-dynamic scatterer component), b, f) *p*_slow_ (fractional contribution of the slow-dynamic scatterer component), c, g) *D*_fast_ (effective diffusion coefficient of fast-dynamic scatterers components), and d, h) *D*_slow_ (effective diffusion coefficient of slow-dynamic scatterers components) derived by bi-exponential model measured during a-d) BH and e-h) HV respiratory challenges. The mean values (solid lines) and corresponding standard deviations (shaded areas) were obtained over 16 datasets measured from eight individuals. The vertical yellow and green regions show the BH and HV periods, respectively.

## 5. Discussion

In this study, we assessed the fractional contribution of static, slow-dynamic, and fast-dynamic scatterers tissue components to the flow index measured by continuous wave diffuse correlation spectroscopy (cw-DCS). To do that, we performed series of data analyses using Siegert relation, modified Siegert relation, and bi-exponential models. The Siegert relation model is the conventional method to derive flow index measured by cw-DCS [24]. In modified Siegert relation model, the contribution of static scatterers can be minimized from the derived flow index. In the bi-exponential model, beside minimizing the contribution of static scatterers, the contribution of slow-dynamic and fast-dynamic scatterer components can be distinguished in the derived flow index. The modified Siegert relation and bi-exponential models are commonly used in data analyses related to laser speckle contrast imaging and interferometric speckle contrast optical spectroscopy [25, 26].

To evaluate the ability of the mentioned three models in assessing the contribution of static, slow-dynamic and fast-dynamic scatterer components to the flow index derived by cw-DCS, two phantoms with controlled medium including Agar-based phantom with tube and liquid phantoms were utilized. Unlike biological tissue with red blood cells (RBCs) [17], nuclei, mitochondria [18], and intracellular components [19], etc., the designed phantoms don’t reflect the complex structure of spatial and depth distributions of tissue components. Instead, these phantoms simplify the complexity of tissue structure and helps to better understand the effects of each medium components to the flow index measured by cw-DCS.

The substantially higher *αD_b_* _(Dynamic)_ (≈ 49%) derived by the modified Siegert relation compared to the *αD_b_* derived by Siegert relation model, measured on Agar-based phantom with tube with relatively high ratio of static scatterers, confirms that the presence of these scatterers leads to underestimation of the cw-DCS derived flow index. This is because the Siegert relation model assumes that all detected photons experience dynamic scattering events and therefore attributes the entire decay of autocorrelation function to dynamic scatterers. While, photons interacting with static scatterers also influence the autocorrelation decay. At baseline, the estimated ratio of light intensity fluctuations caused by static to total scatterer components (*η*) is high (*η* ≈ 1), meaning most photon scattering was originated from the Agar mixture. In response to increased pump flow (*F*_pump_), and increased the contribution of dynamic scatterers in the detected signal, *η* decreased (figure 3c). Importantly, this decrease does not imply changes in number of static scatterers during the experiment. Instead, it reflects the decreased light intensity fluctuations caused by static to total scatterer. The fractional contribution of the fast-dynamic to total scatterers (*p*_fast_) increased substantially by imposing the *F*_pump_ and remained relatively unchanged at all *F*_pump_. As the diameter of the tubes didn’t vary by *F*_pump_, *p*_fast_ remained relatively constant at all *F*_pump_ with some fluctuations in the signal due to the fitting residuals (figure 4 a). These results confirm that *p*_fast_ represents the fraction of fast-dynamic scatterers’ changes. At *F*_pump_ = 0, when there is no fast-dynamic scatterers, the fractional contribution of the slow-dynamic scatterers (*p*_slow_) is the highest. While, by applying the *F*_pump_, the *p*_slow_ substantially decreased and remained relatively constant at all *F*_pump_ (figure 4 b). These results confirm that *p*_slow_ represents the fraction of slow-dynamic scatterers’ changes. As expected, the effective diffusion coefficient of fast-dynamic scatterers (*D*_fast_) derived by bi-exponential model, increased linearly with increasing the *F*_pump_ (figure 4 c), confirming that this parameter primarily represents the displacements of fast-dynamic scatterer components. In contrast, the effective diffusion coefficient of slow-dynamic scatterers (*D*_slow_) remained nearly constant over all *F*_pump_ (figure 4 d), indicating the ability of the bi-exponential model to extract the displacements of slow-dynamic scatterer components. These results are in line with our previous observation measured on the same phantom using the intravoxel incoherent motion (IVIM-MRI) [submitted]. The considerably higher *D*_fast_ compared to *αD_b_* _(Dynamic)_ (≈ 52%) and *αD_b_* (≈ 123%) show the substantial underestimation of the cw-DCS derived flow index due to presence of static scatterers, as well as together, static scatterers and slow-dynamic scatterers of the medium.

The liquid phantoms enabled evaluating the presence of dynamic scatterer components of a medium, to the flow index measured by cw-DCS, while minimizing the presence of static scatterers. In contrast to the Agar-based phantom with tube, where modified Siegert relation model subsequently increased the estimated *aD_b_* _(Dynamic)_, minor differences (17%) were observed between the *aD_b_* and *aD_b_* _(Dynamic)_ measured on the liquid phantoms (figure 5 a, b). Although in a medium with no static scatterers, *η* ≈ 0 is expected theoretically, some photons may undergo relatively fewer scattering events result in a nonzero *η*, causing a deviation from this expectation (figure 5 c). Moreover, the non-zero values of the estimated *η* may originate from residuals of the fitting model rather than presence of true static scatterers. Although the added Glycerol to the liquid phantom decreases the dynamic of scatterers, the substantially higher *p*_fast_ in comparison to *p*_slow_, shows the subsequently higher contribution of fast-dynamic scatterer comparing to slow-dynamic scatterer components to the flow index measured by cw-DCS. The differences in the estimated *p*_fast_ and *p*_slow_ in phantoms with various Glycerol concentration (40% and 20%), confirms that bi-exponential model is not only able to distinguish the contribution of slow-dynamic and fast-dynamic scatterers components to the cw-DCS derived flow index, it is sensitive to the fraction of these scatterers (figure 6 a, b). The relatively low differences (≈ 6%) between the *D*_fast_ and *aD_b_* _(Dynamic)_ shows that slow-dynamic scatterers has minor influence to the cw-DCS derived flow index. Comparing *D*_fast_ and *aD_b_*, indicates the underestimation (≈ 25%) of the flow index derived by Siegert relation, due to presence of both static and slow-dynamic scatterers.

The forearm occlusion test helps to investigate the relative contributions of static, slow-dynamic, and fast-dynamic scatterers to the cw-DCS derived blood flow index (BFI) measured on muscle tissue. The higher (≈ 9%) *BFI*_Dynamic_ in response to forearm occlusion compared to BFI, confirms the underestimation of Siegert relation to estimate the cw-DCS derived blood flow index, which has also been reported in other studies [15, 37, 38]. In response to muscle occlusion, *η* (figure 7 c) and *p*_slow_ (figure 8 b) increased. This observation might be due to the fact that in response to occlusion and decreased the RBCs displacements as fast-dynamic scatterers [17], the ratio of light intensity fluctuations caused by static (extravascular structure of tissue, e.g., nuclei, mitochondria) [18] to total scatterers as well as the fractional contribution of the slow-dynamic scatterers (intracellular motility within the tissue) [19], in the detected signal is increased. As occlusion doesn’t affect the intracellular mobility within the tissue known as slow-dynamic scatterers [19], *D*_slow_ is expected to remain relatively constant. The changes in *D*_slow_ following the occlusion release were comparable to the fluctuations observed throughout the trial (figure 8 d). The higher *D*_fast_ than *BFI*_Dynamic_ (≈ 15%), confirms that presence of slow-dynamic scatterers (intracellular motility within the tissue) [19], led to underestimation of the cw-DCS derived blood flow index. The static scatterers (extravascular structure of tissue, e.g., nuclei, mitochondria) [18], together with slow-dynamic scatterers (intracellular motility within the tissue) [19], cause higher underestimation (≈ 26%) of the cw-DCS derived blood flow index. The hyperemic response which is due to reperfusion following by occlusion release at 180 mmHg was observed in *D*_fast_ (figure 8 c). These results confirm the ability of the bi-exponential model to distinguish the signal decorrelation caused by static, slow-dynamic and fast-dynamic scatterers. The substantial decrease of *p*_fast_ in response to occlusions (figure 8 a), confirms this parameter refers to fractional contribution of RBCs as fast-dynamic scatterers. In our previous study we observed an increase in the IVIM-derived perfusion fraction [Submitted] which resembles the cw-DCS derived *p*_fast_, measured during leg and forearm occlusion test. Venous occlusion leads to decrease the venous outflow while arterial inflow remained unchanged [39, 40]. This causes an increase in tissue blood volume [41] and as a result increased the IVIM-derived perfusion fraction. Thus, the contrast between these results comes from the fact that perfusion fraction derived by IVIM-MRI represent the volume fraction of the blood flow, while, the *p*_fast_ derived by cw-DCS is more sensitive to dynamic of RBCs compared to the volume of the RBCs.

To evaluate the contribution of static, slow-dynamic and fast-dynamic scatterer tissue components on the cerebral blood flow (CBF) changes measured by cw-DCS, breath hold (BH) and Hyperventilation (HV) respiratory challenges were performed. The *BFI*_Dynamic_ is higher than the BFI (3% and 7%) estimated throughout BH and HV respiratory challenges trials. This difference is slightly lower in comparison to the corresponding parameters measured during forearm occlusion test (9%). This might be due to the fact that forearm occlusions, led to more substantial changes in the ratio of light intensity fluctuation caused by static to total scatterers or η (figure 9 c), and as a results higher differences between the *BFI*_Dynamic_ and BFI. However, the η remained nearly constant throughout whole BH and HV respiratory challenge trials. In response to both BH and HV, *p*_fast_ and *p*_slow_ also remained relatively constant (figure 10 a, b). The higher *p*_fast_ than *p*_slow_ confirms substantial higher contribution of RBCs as fast-dynamic scatterers [17] in comparison to slower motility within intracellular structures [19] as slow-dynamic scatterers. The estimated *D*_fast_ was higher than *BFI*_Dynamic_ and BFI throughout both BH (13% and 16%) and HV (15% and 24%) trials. These differences were slightly higher in response to HV than BH, which might be due to physiological effect in response to these respiratory challenges. Although, during baseline, no differences are expected between the cw-DCS derived parameters measured on human head, the minor mismatch between these values may come from the data analyses (e.g. residual in fitting) rather than physiological reasons. The substantially higher *D*_fast_ than *D*_slow_ as well as the relatively constant *D*_slow_ throughout the BH and HV trials (figure 10 c, d) confirms that cw-DCS derived flow index primarily originates from RBCs displacements as fast-dynamic scatterers rather than slower motility within intra-cellular structures [19].

As expected, the η measured on forearm (≈ 0.3) and foreheads (≈ 0.4) were higher than the corresponding parameter measured on liquid phantom (≈ 0.1) and lower than Agar-based phantom with tube (≈ 0.9). As skull is a static scatterer medium, the estimated η measured at baseline on forehead is slightly higher than η measured on forearm. The *p*_slow_ measured on forearm (≈ 0.3) and foreheads (≈ 0.1 − 0.2) were in the range of *p*_slow_ measured on liquid mixture (≈ 0.1) confirming that slow-dynamic scatterer components of tissue have less contribution comparing to fast-dynamic scatterer component in the cw-DCS derived blood flow index. The different η and *p*_slow_ measured on forearm and forehead, show that each tissue type has a specific parameter. Same as laser speckle contrast imaging and interferometric speckle contrast optical spectroscopy [25, 26], the estimation of these parameters derived by cw-DCS, may give useful information about tissues’ structure. However, further improvements in the experimental setup and data-analysis methods are necessary to evaluate their potential for tissue characterization.

Agar-based phantom with tubes was a medium with the most pronounced differences between the dynamic properties of static, slow and fast scatterers. The highest differences between the calculated *αD_b_*_(Dynamic)_ and *αD_b_* (49%) as well as *D*_fast_ and *αD_b_* (123%) on this phantom, showed that modified Siegert relation and bi-exponential models can successfully minimize the influence of static scatterer and slow-dynamic scatterer components on the cw-DCS derived flow index. In case of liquid phantoms, the less differences between the dynamic properties of scatterer components led to lower differences between the *αD_b_*_(Dynamic)_ and *αD_b_* (17%) as well as *D*_fast_ and *αD_b_* (25%). The low differences between the *BFI*_Dynamic_ and BFI (3% - 9%) as well as *D*_fast_ and BFI (13% - 26%) measured on forearm and forehead confirms that, although the tissues extravascular [18], and intracellular mobility within the tissue structure [19] influence the multiple scattering events of photons and decay rate of autocorrelation function, the measured cw-DCS derived BFI is substantially correlated with displacement of RBCs in microvessels [42]. The closer agreement between the obtained results in human tissue and those obtained in the liquid phantoms, compared with the Agar-based phantom with tubes, further supports this interpretation. The *D_fast_* in Agar-based phantom with tubes is two orders of magnitude higher than *D_slow_*. The corresponding ratio in liquid phantom with Glycerol as well as forearm and forehead is an order of magnitude which is in line with the resembling parameters measured by IVIM-MRI [43].

Although the Siegert-relation model underestimates the absolute flow index measured by cw-DCS, the corresponding changes observed during the muscle occlusion and respiratory challenges remained relatively consistent across all three fitting approaches. Thus, the bias introduced by static and slow-dynamic scatterers primarily affects the absolute value of the flow index measured using cw-DCS, derived by Siegert relation, rather than its response to physiological challenges. Nevertheless, when effective diffusion coefficient of fast-dynamic scatterers (*D*_fast_) referring to absolute flow index is required, correction for the static and slow-dynamic scatterer components may improve the accuracy of the cw-DCS derived flow index. However, implementing modified Siegert relation and bi-exponential models come with the cost of a noisier derived parameters.

## 6. Conclusion

In this study, we evaluate the fractional contribution of the static, slow-dynamic, and fast-dynamic scatterer components of the medium to the cw-DCS derived flow index. We performed three data analysis approaches including Siegert relation, modified Siegert relation and bi-exponential models to derive the flow index parameters. Our results confirm that presence of static and slow-dynamic scatterer components lead to underestimation of the absolute flow index measured by cw-DCS derived by Siegert relation. However, the displacements of the red blood cells as fast-dynamic scatterer components dominates the influence of static and slow-dynamic scatterer components in the estimated blood flow index. Thus, employing modified Siegert relation and bi-exponential models are necessary, especially when the absolute blood flow index measured by cw-DCS is required.

## Author contributions

Conceptualization: N.M.; Methodology: N.M., D.B.; Investigation: N.M.; Data curation: N.M.; Formal analysis: N.M.; Software: D.B.; Interpretation of results: N.M., M.K., D.B.; Writing original draft: N.M.; Review and editing: N.M., M.K., D.B.; Supervision: D.B. All authors read and approved the final manuscript.

## Data availability

Data underlying the results presented in this paper may be obtained from the authors upon reasonable request.

## References

1 Ashby N, Squiers J. A Historical Perspective on the Development of Modern Concepts of Tissue Perfusion: Prehistory to the Twentieth Century. Critical Care Nursing Clinics of North America 2014; 26: 297–309.

2 Purkayastha S, Sorond F. Transcranial Doppler ultrasound: technique and application. Seminars in neurology 2012; 32: 411–420.

3 Fredriksson I, Larsson M, Strömberg T. Measurement depth and volume in laser Doppler flowmetry. Microvascular Research 2009; 78: 4–13.

4 Shukla AK, Kumar U. Positron emission tomography: An overview. J Med Phys 2006; 31: 13–21.

5 Liu HL, Pu Y, Liu Y, Nickerson L, Andrews T, Fox PT et al. Cerebral blood flow measurement by dynamic contrast MRI using singular value decomposition with an adaptive threshold. Magn Reson Med 1999; 42: 167–172.

6 O’Sullivan TD, Dehghani H, Re R. Diffuse Optical Spectroscopy: Technology and Applications: introduction to the feature issue. Biomedical Optics Express 2024; 15: 6516–6520.

7 Yu G, Durduran T, Zhou C, Wang H-W, Putt ME, Saunders HM et al. Noninvasive Monitoring of Murine Tumor Blood Flow During and After Photodynamic Therapy Provides Early Assessment of Therapeutic Efficacy. Clinical Cancer Research 2005; 11: 3543–3552.

8 Culver JP, Durduran T, Furuya D, Cheung C, Greenberg JH, Yodh AG. Diffuse optical tomography of cerebral blood flow, oxygenation, and metabolism in rat during focal ischemia. J Cereb Blood Flow Metab 2003; 23: 911–924.

9 Kim MN, Durduran T, Frangos S, Edlow BL, Buckley EM, Moss HE et al. Noninvasive measurement of cerebral blood flow and blood oxygenation using near-infrared and diffuse correlation spectroscopies in critically brain-injured adults. Neurocrit Care 2010; 12: 173–180.

10 Buckley EM, Hance D, Pawlowski T, Lynch J, Wilson FB, Mesquita RC et al. Validation of diffuse correlation spectroscopic measurement of cerebral blood flow using phase-encoded velocity mapping magnetic resonance imaging. J Biomed Opt 2012; 17: 037007.

11 Yu G, Floyd TF, Durduran T, Zhou C, Wang J, Detre JA et al. Validation of diffuse correlation spectroscopy for muscle blood flow with concurrent arterial spin labeled perfusion MRI. Opt Express 2007; 15: 1064–1075.

12 Carp SA, Dai GP, Boas DA, Franceschini MA, Kim YR. Validation of diffuse correlation spectroscopy measurements of rodent cerebral blood flow with simultaneous arterial spin labeling MRI; towards MRI-optical continuous cerebral metabolic monitoring. Biomed Opt Express 2010; 1: 553–565.

13 Shoemaker LN, Samaei S, Deller G, Wang DJJ, Milej D, St Lawrence K. All-optics technique for monitoring absolute cerebral blood flow: validation against magnetic resonance imaging perfusion. Neurophotonics 2024; 11: 045002.

14 Blanco I, Zirak P, Dragojevic T, Castellvi C, Durduran T, Justicia C. Longitudinal, transcranial measurement of functional activation in the rat brain by diffuse correlation spectroscopy. Neurophotonics 2017; 4: 045006.

15 Mogharari N, Kacprzak M, Lipinski K, Liebert A, Wojtkiewicz S. Correlation of Hemodynamic Responses Measured on Human Head by Diffuse Correlation Spectroscopy and BOLD MRI. IEEE Journal of Selected Topics in Quantum Electronics 2025; 31: 1–15.

16 Boas DA, Campbell LE, Yodh AG. Scattering and Imaging with Diffusing Temporal Field Correlations. Physical review letters 1995; 75 9: 1855–1858.

17 Cheung C, Culver JP, Takahashi K, Greenberg JH, Yodh AG. In vivo cerebrovascular measurement combining diffuse near-infrared absorption and correlation spectroscopies. Phys Med Biol 2001; 46: 2053–2065.

18 Ninck M, Untenberger M, Gisler T. Diffusing-wave spectroscopy with dynamic contrast variation: disentangling the effects of blood flow and extravascular tissue shearing on signals from deep tissue. Biomed Opt Express 2010; 1: 1502–1513.

19 Liu B, Postnov D, Boas DA, Cheng X. Dynamic light scattering and laser speckle contrast imaging of the brain: theory of the spatial and temporal statistics of speckle pattern evolution. Biomedical Optics Express 2024; 15: 579–593.

20 Lemieux PA, Durian DJ. Investigating non-Gaussian scattering processes by using nth-order intensity correlation functions. J Opt Soc Am A 1999; 16: 1651–1664.

21 Wu MM, Perdue K, Chan ST, Stephens KA, Deng B, Franceschini MA et al. Complete head cerebral sensitivity mapping for diffuse correlation spectroscopy using subject-specific magnetic resonance imaging models. Biomed Opt Express 2022; 13: 1131–1151.

22 Robinson MB, Renna M, Otic N, Kierul OS, Muldoon A, Franceschini MA et al. Pathlength-selective, interferometric diffuse correlation spectroscopy. IEEE J Sel Top Quantum Electron 2025; 31.

23 Samaei S, Colombo L, Borycki D, Pagliazzi M, Durduran T, Sawosz P et al. Performance assessment of laser sources for time-domain diffuse correlation spectroscopy. Biomed Opt Express 2021; 12: 5351–5367.

24 Durduran T, Choe R, Baker WB, Yodh AG. Diffuse Optics for Tissue Monitoring and Tomography. Rep Prog Phys 2010; 73.

25 Boas DA, Dunn AK. Laser speckle contrast imaging in biomedical optics. J Biomed Opt 2010; 15: 011109.

26 Nowacka-Pieszak K, Samaei S, Borycki D. Interferometric speckle contrast optical spectroscopy (iSCOS) in continuous-wave parallel interferometric near-infrared spectroscopy (CW-πNIRS). Biocybernetics and Biomedical Engineering 2025; 45: 669–684.

27 Bihan DL, Breton E, Lallemand D, Aubin ML, Vignaud J, Laval-Jeantet M. Separation of diffusion and perfusion in intravoxel incoherent motion MR imaging. Radiology 1988; 168: 497–505.

28 Shiraishi T, Chikui T, Inadomi D, Kagawa T, Yoshiura K, Yuasa K. Evaluation of diffusion parameters and T2 values of the masseter muscle during jaw opening, clenching, and rest. Acta Radiologica 2012; 53: 81–86.

29 Becker W. Advanced Time-Correlated Single Photon Counting Applications, 2005.

30 Sudakou A, Wabnitz H, Liemert A, Wolf M, Liebert A. Two-layered blood-lipid phantom and method to determine absorption and oxygenation employing changes in moments of DTOFs. Biomed Opt Express 2023; 14: 3506–3531.

31 Cortese L, Presti GL, Pagliazzi M, Contini D, Mora AD, Pifferi A et al. Liquid phantoms for near-infrared and diffuse correlation spectroscopies with tunable optical and dynamic properties. Biomed Opt Express 2018; 9: 2068–2080.

32 Irwin D, Dong L, Shang Y, Cheng R, Kudrimoti M, Stevens SD et al. Influences of tissue absorption and scattering on diffuse correlation spectroscopy blood flow measurements. Biomed Opt Express 2011; 2: 1969–1985.

33 Liebert A, Wabnitz H, Grosenick D, Möller M, Macdonald R, Rinneberg H. Evaluation of optical properties of highly scattering media by moments of distributions of times of flight of photons. Appl Opt 2003; 42: 5785–5792.

34 Zhu Y, Gui Z, Xue B, Shang Y. Experimental Validation of Microvasculature Blood Flow Modeling by Diffuse Correlation Spectroscopy. IEEE Access 2020; 8: 15945–15951.

35 Durduran T, Yu G, Burnett MG, Detre JA, Greenberg JH, Wang J et al. Diffuse optical measurement of blood flow, blood oxygenation, and metabolism in a human brain during sensorimotor cortex activation. Opt Lett 2004; 29: 1766–1768.

36 Mogharari N, Wojtkiewicz S, Borycki D, Liebert A, Kacprzak M. Time-domain diffuse correlation spectroscopy at large source detector separation for cerebral blood flow recovery. Biomedical Optics Express 2024; 15: 4330–4344.

37 Bartlett MF, Oneglia AP, Ricard MD, Siddiqui A, Englund EK, Buckley EM et al. DCS blood flow index underestimates skeletal muscle perfusion in vivo: rationale and early evidence for the NIRS-DCS perfusion index. J Biomed Opt 2024; 29: 020501.

38 Durduran T, Minkoff DL, Kim MN, Hance D, Buckley EM, Tobita M et al. Concurrent MRI and Diffuse Correlation & Near-Infrared Spectroscopic Measurement of Cerebral Hemodynamic Response to Hypercapnia and Hyperoxia. Biomedical Optics and 3-D Imaging. Optica Publishing Group: Miami, Florida, 2010, p BTuB2.

39 Loenneke JP, Pujol TJ. The Use of Occlusion Training to Produce Muscle Hypertrophy. Strength & Conditioning Journal 2009; 31: 77–84.

40 Junejo RT, Ray CJ, Marshall JM. Cuff inflation time significantly affects blood flow recorded with venous occlusion plethysmography. Eur J Appl Physiol 2019; 119: 665–674.

41 Kamshilin AA, Zaytsev VV, Mamontov OV. Novel contactless approach for assessment of venous occlusion plethysmography by video recordings at the green illumination. Scientific Reports 2017; 7: 464.

42 Carp SA, Roche-Labarbe N, Franceschini MA, Srinivasan VJ, Sakadžić S, Boas DA. Due to intravascular multiple sequential scattering, Diffuse Correlation Spectroscopy of tissue primarily measures relative red blood cell motion within vessels. Biomed Opt Express 2011; 2: 2047–2054.

43 Le Bihan D. Intravoxel incoherent motion perfusion MR imaging: a wake-up call. Radiology 2008; 249: 748–752.

